# The persistence of locally adapted polymorphisms under mutation swamping

**DOI:** 10.1101/2024.06.18.599577

**Authors:** Takahiro Sakamoto, James R. Whiting, Samuel Yeaman

## Abstract

Locally adapted traits can exhibit a wide range of genetic architectures, from pronounced divergence at a few loci to small allele frequency shifts at many loci. The type of architecture that evolves depends strongly on migration rate, as weakly selected loci experience swamping and do not make stable contributions to divergence. Simulations from previous studies showed that even when mutations are strongly selected and should resist migration swamping, the architecture of adaptation can collapse and become transient at high mutation rates. Here, we use an analytical two-population model to study how this “mutation swamping” phenomenon depends upon population size, strength of selection, and parameters determining mutation effects. To do this, we developed a mathematical theory based on the diffusion approximation to predict the threshold mutation rate above which swamping occurs, and find that this performs well across wide range of parameter space, based on comparisons with individual-based simulations. The mutation swamping threshold depends most strongly on the average effect size of mutations, and weakly on the strength of selection, but is only minimally affected by population size. Across a wide range of parameter space, we observe that mutation swamping occurs when the trait-wide mutation rate is 10^*−*3^–10^*−*2^, suggesting that this phenomenon is potentially relevant to complex traits with a large mutational target. On the other hand, based on the apparent stability of genetic architecture in many classic examples of local adaptation, our theory suggests that per-trait mutation rates are often relatively low.

## Introduction

Understanding genome evolution underlying adaptation is a major goal of evolutionary biology. Local adaptation is a common way for species to adapt to spatially varying environments, where phenotypic divergence caused by genetic change confers a local fitness advantage. Despite ample evidence at the phenotypic level (Felsenstein 1976; Leimu and Fischer 2008; Hereford 2009; Wadgymar *et al*. 2022), until recently, identifying its causal loci has been challenging because most traits have a polygenic basis with causal loci distributed across the genome. However, the availability of large-scale genomic data is changing this situation, and it is increasingly possible to identify the genome-wide basis of local adaptation. Recent studies have found diverse patterns of genetic architectures underlying local adaptation (Nosil *et al*. 2009; McKown *et al*. 2014; Bomblies and Peichel 2022). Phenotypic variation of some traits is largely explained by a small number of genomic regions (Tavares *et al*. 2018), indicating the presence of large effect alleles or tightly linked clusters of small effect alleles. In interesting cases, large effect clusters form in a haplotype block with structural variations (Todesco *et al*. 2020; Hager *et al*. 2022), indicating that tight linkage was beneficial in the course of local adaptation. In other traits, causal alleles are scattered throughout the genome, and a phenotypic difference arises from the collective effect of many loosely linked loci (Daub *et al*. 2013; White *et al*. 2015; Lind *et al*. 2018; Barghi *et al*. 2019; Kess and Boulding 2019). Although concrete evidence for this “diffuse” case is still scarce because of insufficient statistical power, given the highly polygenic basis of many traits, the diffuse architecture might be widespread (Savolainen *et al*. 2013).

Despite the diversity of genetic architecture, it is still unclear why a concentration of causal alleles evolves in some cases and not in others. Previous theoretical studies have investigated this problem, especially focusing on the role of migration-selection balance (Yeaman and Whitlock 2011; Yeaman 2015; Rafajlović *et al*. 2016). One clear pattern is that migration promotes the evolution of a concentrated architecture when the mutation rate is not high. With migration, large effect alleles under strong selection resist migration and maintain their divergence, while small effect alleles tend to be swamped. However, in the vicinity of a large effect allele, the effective migration rate is reduced by linked selection (Bengtsson 1985; Barton and Bengtsson 1986), which enables establishment of small effect alleles (Aeschbacher and Bürger 2014; Yeaman *et al*. 2016) and prevents the loss of established alleles (Aeschbacher and Bürger 2014; Rafajlović *et al*. 2016). As time goes on, many small effect alleles accumulate in linkage with a large effect allele while swamped in other regions, forming a concentrated architecture.

In contrast to the well-established impact of migration, less attention has been paid to the effect of mutation rate. A potentially important but somewhat overlooked pattern is that, when the mutation rate is high, a concentrated architecture does not evolve, even when migration is sufficiently weak relative to selection to prevent migration swamping of a cluster of causal alleles (Yeaman 2015, 2022). In this region of parameter space, local adaptation is caused by the collective effect of many small effect mutations, none of which maintain divergence for a long time. The genetic architecture is highly transient with fast replacement of causal alleles under the balance of frequent mutations and migration swamping. This pattern was observed in simulations with a mutation rate of *µ*∼ 10^*−*3^ or 10^*−*2^ per trait, so it is potentially relevant to complex traits that are predicted to have a particularly large mutational target (∼1 Mb) (Sella and Barton 2019). However, as there is no comprehensive analytical theory so far, fundamental questions remain unanswered: (i) what is the threshold mutation rate that causes a concentrated architecture to transition into a transient one, (ii) how does the threshold change as various parameters change, and (iii) why does the concentrated architecture become unstable above the threshold?

In this study, we present a mathematical theory to describe the transition of a genetic architecture from stable and clustered to transient and diffuse and answer these questions. Since the transition involves both a clustered large effect locus and scattered small effect loci, we developed a new analytical framework in which both types of loci evolve in a coordinated manner. This approach contrasts with previous theories of local adaptation, which considered the two types of loci in separate models: large effect alleles have been considered in the population genetic models (Haldane 1930; Wright 1931; Pollak 1966; Barton 1987; Yeaman and Otto 2011; Aeschbacher and Bürger 2014; Tomasini and Peischl 2018; Sakamoto and Innan 2019), while the collective effect of small effect loci has been analyzed in the quantitative genetics framework (Hendry *et al*. 2001; Débarre *et al*. 2015). Since neither framework can consider their interaction, we need to construct a new model to represent their combined evolution underlying the transition of a genetic architecture. It should be noted that Dekens *et al*. (2022) recently combines the adaptive dynamics of a quantitative trait with a one locus two allele model to discuss the eco-evolutionary dynamics in the local adaptation process. However, since their model assumes very small genetic variation for the quantitative trait, it cannot be applied to our situation in which ample genetic variation exists.

In the following, we first describe our model in which a focal locus of large effect and many background loci of small effect jointly drive trait evolution and show how different genetic architectures evolve in our model. Next, we present a mathematical analysis to derive the threshold mutation rate at which the transition from a concentrated architecture to a transient one occurs. We demonstrate that the transition occurs because a high mutation rate increases genetic variation contributed by small effect loci and promotes phenotypic adaptation, which reduces the selective advantage of the large effect locus in local adaptation, resulting in collapse of its polymorphism. Given the similarity to the swamping that occurs when migration is high relative to selection, we term this effect “mutation swamping”. We compared our theory with the simulation results and confirmed the validity of our analysis. We also investigated how the threshold mutation rate depends on various parameters, including population size, migration rate, and selection strength. The present study provides a fundamental theory on what type of genetic architecture evolves for traits with large mutational targets.

## Model

We use a discrete generation model with two diploid populations (populations 1 and 2), where the size of each population is *N*_1_ and *N*_2_, respectively (Figure 1). Divergent selection acts on a phenotypic trait, whose optimum differs in the two populations. The fitness of an individual with a phenotypic value *z* in population *i* is given by:

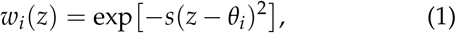

where *s* is the strength of selection, and *θ*_*i*_ is the optimal phenotypic value in population *i*. Throughout this paper, we assume *θ*_1_ = 0 and *θ*_2_ = 1. Migration is assumed to be unidirectional from population 1 to population 2 with rate *m*, following the continent-island model. The effects of selection and migration are incorporated in a reproduction event. In population 1, each parent is chosen from individuals within population 1 with probability weighted by fitness *w*_1_. In population 2, with probability *m*, a parent is chosen from population 1, and otherwise chosen from population 2, based on the probability weighted by fitness.

**Figure 1.**
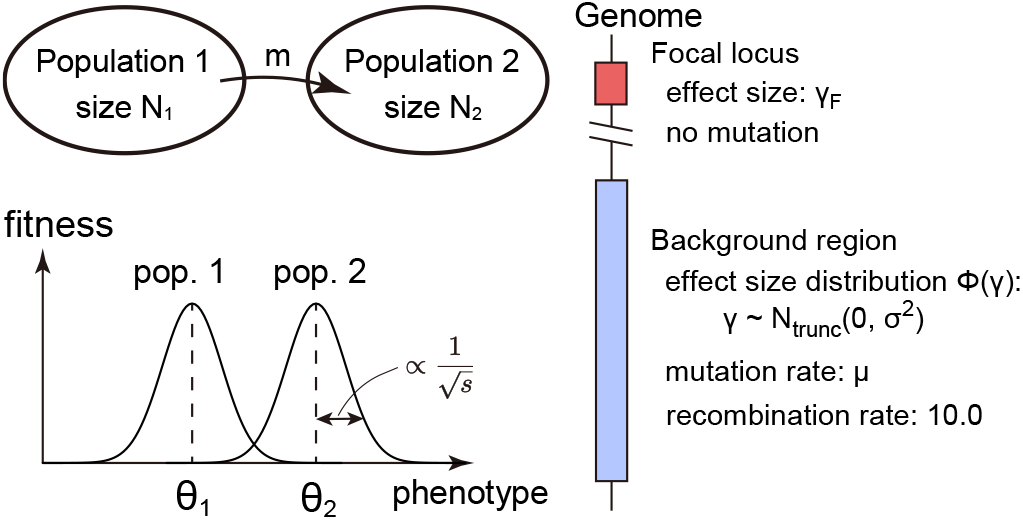
Illustration of the two-population model. See also Table S1 for the list of parameters.

Each genome consists of a focal locus of large effect (which can be interpreted as a single allele of large effect or many alleles of small effect with complete linkage) and a background region that includes many loci of small effect. Phenotype is determined by the additive effect of all causal alleles across both regions. The focal locus has an ancestral allele with an effect size of 0 and a derived allele with an effect size of *γ*_*F*_ (0 *<* 2*γ*_*F*_ *<* 1). During evolution, no mutations arise at the focal locus while mutation occurs in the background region at an overall rate *µ*. In simulations, the background region is subdivided into 1,000 loci, and new mutations are assigned to one of them at random. The effect size of a new mutation is drawn from a truncated normal distribution:

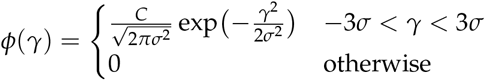

where *C≈* 1.003. Truncation is introduced so that no large effect allele arises at the background region. When a mutation of effect size *γ* arises at a locus with a genotypic value *γ*_0_, the genotypic value of that locus after the mutation becomes *γ*_0_ + *γ*.

The number of recombination events within the background region is determined by the Poisson distribution with a mean of 10, which follows a typical order of a genome-wide recombination rate (Kong *et al*. 2002; Brazier and Glémin 2022). Free recombination is assumed between the focal locus and the background region.

Since we are interested in the stability of a focal locus of large effect, we start evolution from a state in which the focal locus is divergent. At the focal locus, the ancestral allele is fixed in population 1 while the derived allele is fixed in population 2. In the background region, a genotype with an effect size of 0 is fixed in the entire population, so that an initial phenotypic value in each population is 0 and 2*γ*_*F*_, respectively. In the simulations, starting from this state, we ran simulations for 200,000 generations and registered the population state every 1,000 generations. We changed various parameters including migration and mutation rates and investigated their effect on the genetic architecture. We ran five replicates for each parameter set.

## Results

### Overview of observed genetic architecture

From extensive simulations, we rediscover the three patterns of genetic architecture (Figure 2A–C), consistent with Yeaman (2015, 2022): a diffuse architecture observed when *m* is small, a concentrated architecture observed when *m* is moderate and *µ* is small, and a transient architecture observed when *m* is moderate and *µ* is large (Figure 2D). As we mentioned in the Introduction, the present study aims to clarify the role of *µ* in the transition from a concentrated architecture to a transient architecture. However, before moving into the analysis, it is helpful to define how we assess these differences among types of architecture. In this section, we briefly overview the characteristics of each genetic architecture with reference to typical simulation runs.

**Figure 2.**
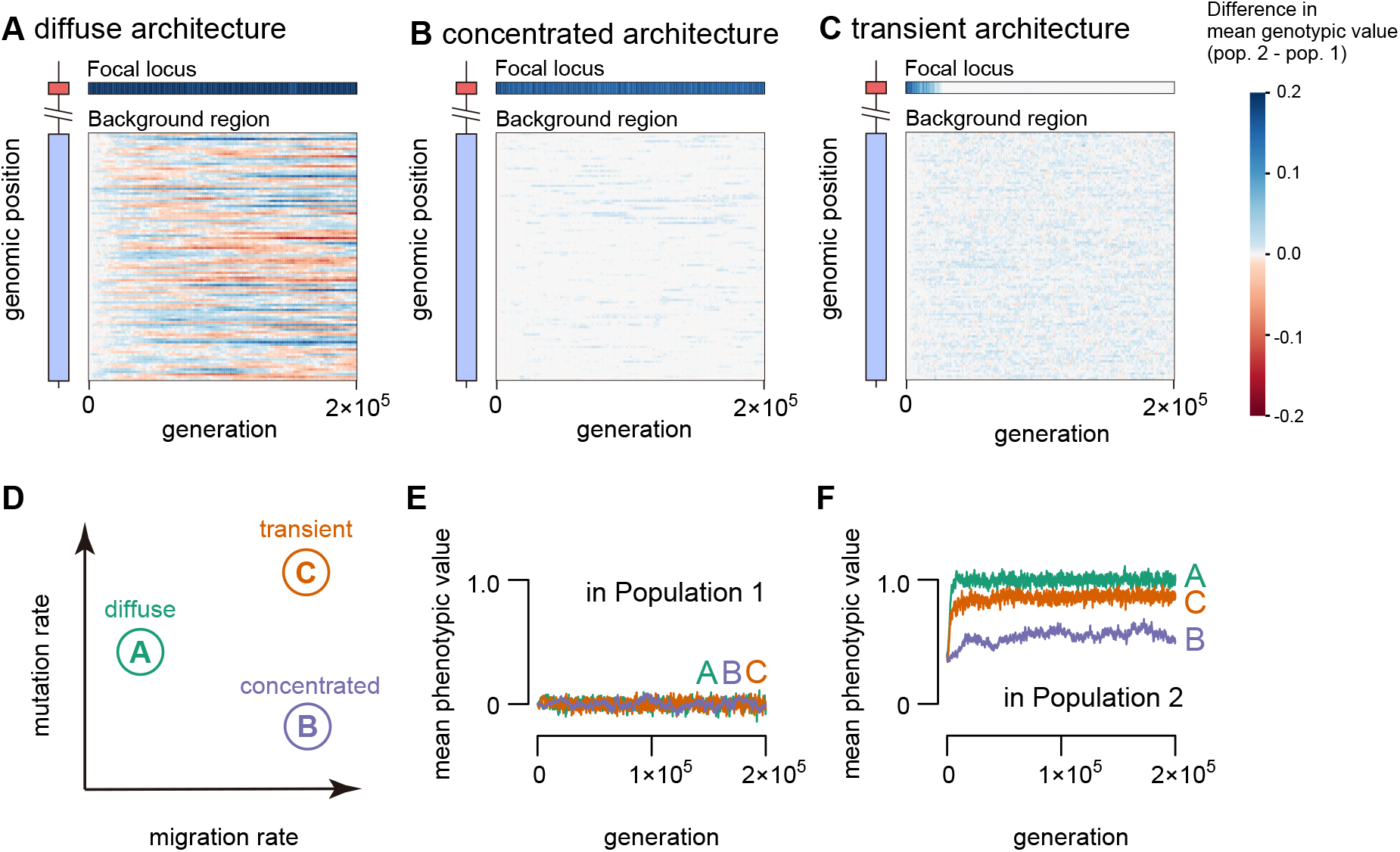
Evolution of genetic architecture in three representative cases of simulation. (A–C) Evolution of effect size divergence between the two populations at the focal locus (top row) and in the background region (bottom subpanel). Following migration and mutation rates were assumed in each panel: (A) *m* = 10^*−*5^ and *µ* = 10^*−*2.25^, (B) *m* = 10^*−*3^ and *µ* = 10^*−*3.25^, and (C) *m* = 10^*−*3^ and *µ* = 10^*−*2^. For other parameters, *N*_1_ = *N*_2_ = 10, 000, *s* = 0.025, *σ* = 0.02, *γ*_*F*_ = 0.2 were assumed (see also Table S1). For visibility, the background region is composed into 100 rows, each row representing the sum of the genotypic value difference of 10 adjacent loci. (D) A schematic diagram for the migration and mutation rates assumed in the three cases. (E, F) Mean phenotypic value in population 1(E) and population 2 (F) in the three cases.

A diffuse architecture refers to a situation in which highly divergent loci are distributed across the genome. This architecture is observed when *m* is so small that small effect alleles are able to resist migration. After many generations, the focal locus remains divergent, while many small effect mutations accumulate in the background region, constituting a heterogeneous landscape of effect sizes (Figure 2A). Some regions evolved effect sizes comparable to the focal locus, resulting in many large effect loci diffused throughout the genome. The phenotypic value is very close to the optimum in both populations, and local adaptation is almost maximized (Figure 2E, F).

In a concentrated architecture, the divergence is limited to the focal locus, and no stable divergence is observed in the background region (Figure 2B). At the focal locus, the divergence is maintained at migration-selection balance, explaining a large part of the phenotypic difference between the populations. Migration works to prevent the divergence of small effect alleles in the background region, resulting in intermediate phenotypic divergence (Figure 2E, F).

A transient architecture is a state where no stable divergence is maintained at any loci in the genome (Figure 2C). This state is observed at a higher mutation rate than the rate at which the concentrated architecture is observed. The focal locus is no longer able to maintain its divergence and is homogenized by migration. Although the divergence at small effect loci is still inhibited, local adaptation is pronounced at the phenotypic level (Figure 2E, F), indicating the role of the cumulative effect of small frequency differences across many loci. Figure 2C shows that the causal loci underlying local adaptation turnover frequently, demonstrating that the genetic architecture is highly transient.

In the following section, we specifically focus on the transition from a concentrated architecture to a transient architecture. First, we provide a theoretical framework to explain why this transition occurs and how the threshold mutation rate is determined. Next, we compare the theoretical prediction with the simulation results and see how various parameters affect the threshold.

### Theory on the transition between concentrated architecture and transient architecture

To seek the reason why the transition from a concentrated architecture to a transient one occurs as the mutation rate increases, we perform a mathematical analysis of the model. The most pronounced difference between the two architectures is the persistence time of divergence at the focal locus (see Figure 2B, C). Thus, in this section, we specifically focus on the divergence at the focal locus in addition to the phenotypic divergence. Let 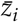 and *p*_*i*_ be the mean phenotypic value and the derived allele frequency at the focal locus in population *i*, respectively. These quantities in population 1 are straightforward: Under the unidirectional migration, it evolves independently from population 2 to 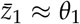 and *p*_1_ = 0. Thus, our aim in this section is to derive the formula for 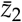 and *p*_2_. If *p*_2_ = 0 is the only stable equilibrium, we predict that the concentrated architecture is unstable and the transient architecture evolves.

Our analysis takes the following steps. First, we suppose that 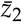 reaches its equilibrium value. Then, we can calculate *p*_2_ and mean genotypic value for the background region, 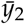, as a function of 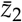. If 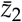 is the equilibrium, it should satisfy the self-consistency condition:

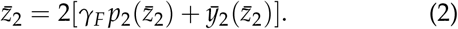

Using this condition, we determine 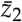 and *p*_2_ numerically. Throughout this analysis, we assume that recombination works efficiently and the linkage effect can be ignored.

### Dynamics at the focal locus

Under the linkage equilibrium approximation, the dynamics of *p*_2_ are described by:

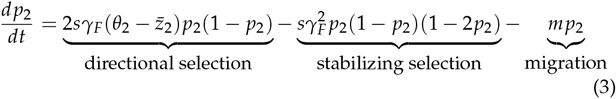

(see Barton 1986; de Vladar and Barton 2014; Jain and Stephan 2015). In Equation 3, the first term and the second term arise due to directional and stabilizing selection on the phenotypic value, respectively. A required condition for Equation 3 to have a stable equilibrium 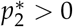 is given by:

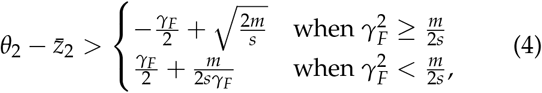

and, if 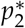 exists, it is calculated as:

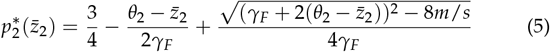

(see APPENDIX A for details).

Equation 4 explains why the concentrated architecture does not evolve at a high mutation rate. This equation shows that the divergence at the focal locus is unstable when 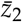 is too close to *θ*_2_ because selection on the derived allele is weak when the phenotype is well adapted (see the directional selection term in Equation 3). Once selection becomes weaker than migration, the derived allele at the focal locus is swamped by migration. Since mutation increases the genetic variation at the background region and fuels adaptation (Hendry *et al*. 2001), it could lead to the collapse of its divergence.

One important prediction of Equation 4 is that this transition behavior along the mutation rate is observed only when the migration rate is within a certain range.

When 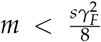, a stable equilibrium exists even with maximal phenotypic divergence (i.e.,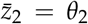). In this regime, migration is much weaker than disruptive selection arising from stabilizing selection on phenotype, which is represented by the second term in the right hand of Equation 3. Then, starting from the divergent state, its divergence is stably maintained. In the opposite limit of 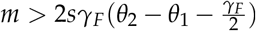, no stable equilibrium exists even in the maximum possible maladaptation (i.e.,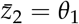). In this parameter range, migration always overwhelms selection and inhibits divergence. Between these two migration rates, stability of the divergence at the focal locus depends on the extent of phenotypic adaptation. As the mutation rate increases, genetic variation at the background region increases, and the phenotypic value becomes closer to its optimum. Then, the divergence at the focal locus becomes unstable. Below, we derive the threshold of the mutation rate through explicitly quantifying the phenotypic adaptation driven by the background region.

### Dynamics at the background region

To predict the threshold mutation rate, we need to calculate 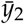. Under the linkage equilibrium approximation, 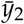 is determined by the additive effects of all segregating alleles. To quantify it, we first consider the behavior of a mutant allele. Next, we sum up the contribution of each mutant allele and derive 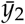. Since a new mutant allele can arise in either population, we consider these cases separately.

In the derivation, an important approximation we made is that the effect of fixation of a mutant allele occurring in population 1 is small. This approximation may be justified because fixation should be rare at equilibrium. Under stabilizing selection on the phenotype, disruptive selection works on each mutant allele (Kimura 1981, see also Equation 3), preventing its increase. Even when a fixation occurs, the fixed allele rapidly spreads in population 2 in the face of gene flow and does not contribute to phenotypic divergence significantly. Below, we consider only mutant alleles that arise in population 2 or alleles that arise in population 1 and go extinct.

First, we consider a mutant allele that arises in population 2 with an effect size *γ* larger than the ancestral allele. Since the ancestral allele is fixed in population 1, under unidirectional migration, the long-term fate of this allele is extinction. The total contribution of this allele to 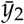 before its extinction is given by

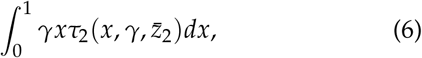

where 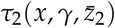 is the sojourn time density at frequency *x* in population 2. In APPENDIX B, we derived the explicit form of 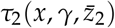 based on diffusion theory.

Next, we consider a mutant allele that arises in population 1, which can contribute to 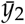 when it migrates to the population 2 and stays there for some time. We first consider the dynamics in the population 1. Assuming its long-term extinction, the expected number of alleles that migrate to population 2, *n*_1_(*γ*), is given by:

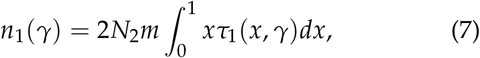

where *τ*_1_(*x, γ*) is a sojourn time density of the mutant allele at frequency *x* in population 1 conditional on its eventual extinction (see APPENDIX B for details). After migration, each mutant allele has a frequency 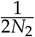, the same as a mutant allele that arises in population 2. Thus, each migrant allele contributes to 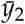 by 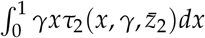.

Now, we calculate 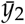 from the total contribution of all mutant alleles that arise in a generation. The expected number of new mutations in population *i* is given by 2*N*_*i*_*µ*. Since the genotypic value of the background region having no mutant allele is 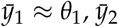 is calculated as:

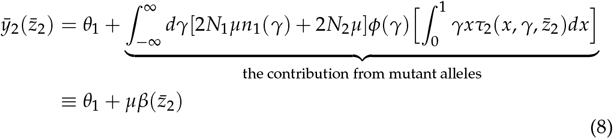

We now have *p*_2_ and 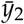 as a function of 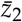, so we can determine the equilibrium values of 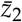 from the self-consistency condition (Equation 2). If there is an equilibrium with *p*_2_ *>* 0, an equilibrium with divergence at the focal locus exists.

### Threshold mutation rate

The threshold mutation rate can be obtained from Equations 4 and 8. Let us assume that 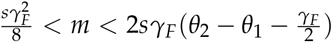 so that the threshold exists. At the threshold, the left hand and the right hand of Equation 4 should be equal. Thus, we can derive:

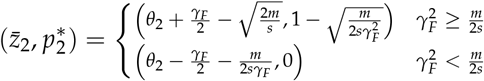

From Equation 2, 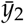 is calculated as:

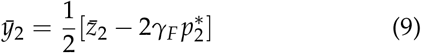

Using Equation 8, the threshold mutation rate is given by:

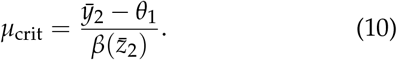

In the following section, we evaluate the performance of Equation 10 by comparing its prediction with the simulation results.

### Simulation results

To check the agreement between predicted threshold mutation rate and observed genetic architecture, we ran individual-based simulations. To distinguish the different architectures based on the simulation output, one may consider a simple indicator based on the derived allele frequency at the focal locus in population 2, which should be > 0 if a concentrated architecture is stable, and otherwise, should be 0. This indicator works in many cases, however, we found that in some cases, the architecture is still concentrated while the divergence at the focal locus disappears (see Figure S1 for an example). In these runs, a large effect locus arises in the background region by tight linkage of small alleles and takes over the role of the cluster. Thus, we need an indicator that is robust to these dynamics.

To determine the architecture more accurately, we developed another indicator that focuses on the stability of the genetic architecture. In both diffuse and concentrated architectures, the same loci tend to contribute to divergence for a long time, whereas in a transient architecture, the causal loci driving divergence turnover frequently (Figure 2B, C). Based on this intuition, we first identified the loci that significantly contributed to the phenotypic divergence at each time point and checked whether these loci were stable or replaced along the time points. We used the simulation output from 101 timepoints between generation 100,000 and 200,000, which is registered every 1,000 generations. The output from the initial 100,000 generations is ignored since we are interested in the equilibrium state. At each time point, we compare the divergence of the genotypic value at each locus (including the focal locus), and identify the loci whose divergence is within the top 1% among all loci. Then, for each locus we calculate the maximum duration during which the locus stays a member of the top loci, *T* (0 ≤ *T* ≤ 101), and took *T*^*∗*^ = max(*T*)/101 as an indicator of the stability of a genetic architecture. This indicator is more robust to the turnover of the cluster because the new cluster locus generally had a large effect size long before the turnover, having a large *T*.

To visualize the threshold mutation rate, we plotted *T*^*∗*^ while changing the migration rate and the mutation rate (Figure 3A). We can see the existence of the threshold, at which the stability of the architecture (*T*^*∗*^) changes significantly. The black line shows the theoretical prediction (Equation 10), which generally agrees well with the threshold observed in the simulations. As expected by Equation 4, the transition is observed over a limited range of migration rates, and a transient architecture evolves at very high migration rate (i.e., migration swamping).

**Figure 3.**
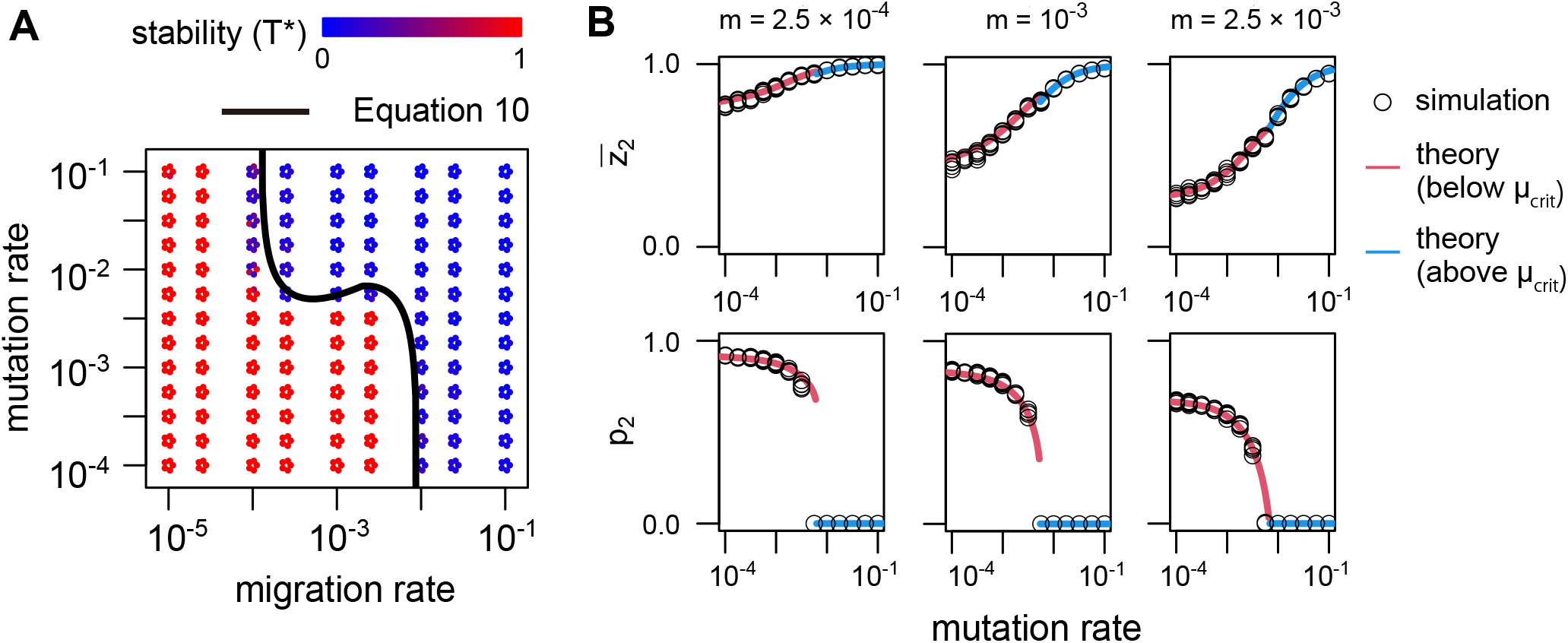
Transition of genetic architecture in simulation. (A) Stability of genetic architecture (*T*^*∗*^) in each parameter set. Each circle shows *T*^*∗*^ calculated from one simulation run, where red color means stable architecture while blue color means unstable one. For each parameter set, five replicates were run. (B) Phenotypic divergence and frequency of the mutant allele at the focal locus as a function of mutation rate. Parameters *N*_1_ = *N*_2_ = 10, 000, *s* = 0.025, *σ* = 0.02, and *γ*_*F*_ = 0.2 were assumed. Solid lines show theoretical prediction.

To examine the mechanism of the transition, we focus on the divergence at the phenotypic level (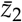) and at the focal locus (*p*_2_). Figure 3B shows how 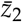 and *p*_2_ change as the mutation rate increases for the three migration rates in which the theory predicted the transition. Each circle represents the average value in the latter half of one simulation, while solid lines are the theoretical predictions calculated from Equations 2, 5, and 8. We observe a good agreement between the theory and the simulation results. In all cases, as the mutation rate increases, 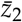 increases because the supply of genetic variation by mutation promotes local adaptation (Hendry *et al*. 2001). Conversely, the divergence at the focal locus (*p*_2_) decreases as the mutation rate increases. When the mutation rate is low, phenotypic adaptation is insufficient and the divergence at the large effect locus is highly advantageous. However, when the mutation rate is high and the phenotype is already close to the optimum, such a large effect allele is no longer necessary, and selective advantage diminishes (see the directional selection term in Equation 3). Therefore, selection at the focal locus decreases as the mutation rate increases, and the divergence becomes unstable when selection becomes weaker than migration.

An interesting pattern in Figure 3 is that the threshold mutation rate is roughly constant within the range of the migration rate where the transition occurs. This pattern is explained by the two effects of migration. While migration reduces divergence at the focal locus through its homogenizing effect, it simultaneously inhibits the divergence at the phenotypic level (Figure 3B) and makes the divergence of the focal locus more advantageous. These effects seem to be well balanced and cancel out the effect of migration on the threshold.

In Figure 3A, we can see some discrepancies between the theory and the simulation results when *m* = 10^*−*4^. It may be because it is near the threshold and the divergence at the focal locus is not too stable. In this region of parameter space, *T*^*∗*^ tends to take intermediate values, supporting our reasoning.

### Effect of various parameters on the threshold

In this section, we investigate the effect of each parameter on the threshold mutation rate (see Table S1 for the list of parameters). To see the overall trend, we calculate the threshold mutation rate from Equation 10 while changing various parameters. In the following, we discuss the effect of each parameter based on the theoretical prediction, which allows the clearest and most general exploration of factors affecting the threshold, but note that we also performed simulations and confirmed the consistency between the theory and the simulation results in these parameters (see Figures S2–S5).

First, we investigated the effect of population size by studying cases where either the island or the continent had a larger population, or where both patches had symmetrical population size. Figure 4A shows that a larger size of population 1 (i.e., continent population) decreases the threshold, whereas changing the size of population 2 (i.e., island population) has no detectable effect. This difference reflects the difference of the evolutionary forces acting on small effect alleles in each population. In population 1, the phenotypic value is close to its optimum and selection pressure is weak, especially on small effect alleles. Thus, evolution is determined by mutation-selection-drift balance. As the size of population 1 increases, genetic variation in population 1 increases, and migrants bring more variation into population 2. Therefore, a lower mutation rate is needed to cause the transition to a transient architecture. In contrast, population 2 is faced with moderate migration in the parameter space where the threshold occurs. Under these conditions, random genetic drift is too weak compared with migration, and the dynamics are mainly determined by mutation-selection-migration balance. Since all effective forces are the deterministic ones, the size of population 2 does not affect the threshold rate very much.

**Figure 4.**
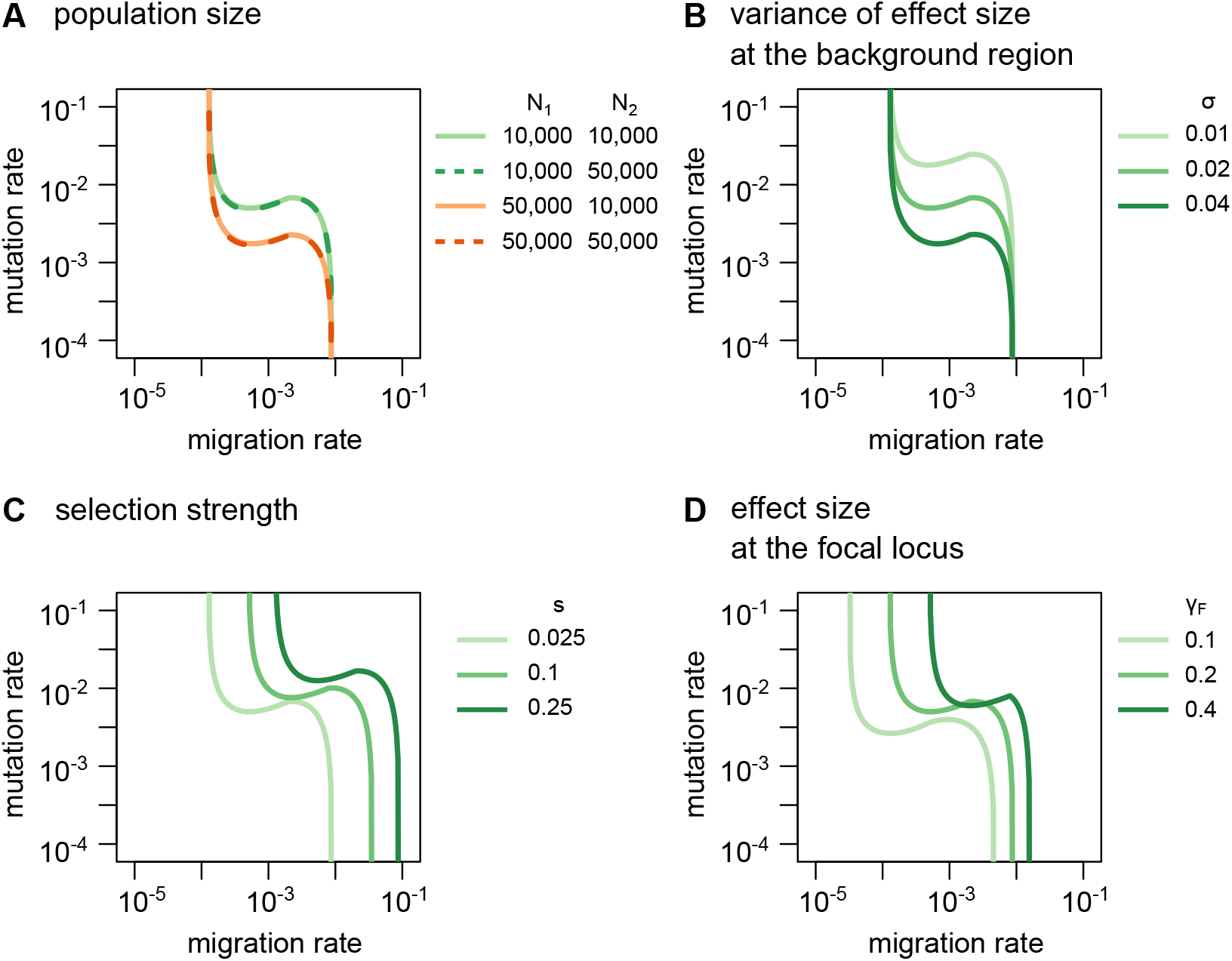
Effect of each parameter on the threshold mutation rate at which the transition of genetic architecture occurs. Unless otherwise specified, *N*_1_ = *N*_2_ = 10, 000, *s* = 0.025, *σ* = 0.02, and *γ*_*F*_ = 0.2 were assumed.

As the variance of the effect size of new mutations in the genetic background (*σ*) increases, the threshold mutation rate decreases (Figure 4B). Since larger *σ* increases the amount of phenotypic variation, it promotes phenotypic adaptation and causes the transition at a lower mutation rate.

We examined the effect of selection strength (*s*) in Figure 4C. As selection becomes stronger, the range of migration rates within which the transition occurs shifts towards higher values, because more migration is needed to overwhelm selection. The threshold mutation rate slightly increases as selection increases because the divergence at the focal locus is more stable under strong selection; Equation 4 shows that the divergence at the focal locus becomes more robust to phenotypic adaptation as selection increases. The effect of selection strength on the threshold is relatively small because stronger selection also enhances adaptation by small effect variants and promotes the phenotypic adaptation.

The impact of the effect size at the focal locus (*γ*_*F*_) is shown in Figure 4D. Similar to the effect of the selection strength, larger 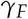 increases the stability of divergence and shifts the migration range where the transition occurs to higher rates. The threshold mutation rate tends to increase as *γ*_*F*_ increases, but its effect seems relatively small, especially for large *γ*_*F*_. This pattern arises through a complex interaction between the focal locus and the background region. Although the divergence at the focal locus becomes more robust to phenotypic optimization as *γ*_*F*_ increases (Equation 4), larger *γ*_*F*_ also increases the relative contribution of the focal locus to phenotypic divergence.

Of note, if we calculate the amount of divergence in the background region necessary to cause the transition using Equation 9, we find the threshold 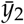 decreases as *γ*_*F*_ increases when 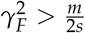. Thus, despite more stability at the focal locus, smaller divergence in the background region is enough to cause the transition. However, simultaneously, larger effect of the focal locus promotes phenotypic divergence, which reduces the selection strength acting on small effect variants. Then, more mutations are needed to yield a given amount of divergence in the genotypic value driven by the background region. These counteracting effects of *γ*_*F*_ result in its relatively small impact on the threshold mutation rate when *γ*_*F*_ is large.

Although we have so far assumed a unidirectional migration for mathematical simplicity, a bidirectional migration between subpopulations would be another common situation. For completeness, we also used a symmetric two-population model, in which migration from population 2 to population 1 was allowed at rate *m* and *N*_1_ = *N*_2_ = *N* was assumed. Using this model, we checked the generality of the qualitative patterns we presented in this section. Since our mathematical approximations are inapplicable to this situation, we ran individual-based simulations to investigate the transition of genetic architecture. Figure S6 shows a qualitatively very similar pattern as Figure 4, indicating that our theoretical arguments above are also applicable to the case of bidirectional migration.

## Discussion

How phenotypic divergence underlying local adaptation is encoded in the genome has been a major focus of population genetics. Species in nature exhibit a variety of genetic architectures ranging from monogenic, oligogenic, to polygenic architecture (Nosil *et al*. 2009; McKown *et al*. 2014; Bomblies and Peichel 2022). However, the evolutionary factors that determine genetic architecture remain unclear. The present study highlights the role of mutation rate at loci determining a focal phenotype. When the mutation rate is low, a concentrated architecture with a small number of diverged large effect loci evolves in the face of the gene flow. However, as the mutation rate increases, divergence at large effect loci becomes unstable, and the architecture transitions to a transient one driven by fleeting divergence at many small effect loci dispersed across the genome. We termed this collapse of the stable divergent architecture “mutation swamping”, with an analogy to the classic migration swamping (Haldane 1930; Wright 1931; Felsenstein 1976; Lenormand 2002). Analogous behaviour with the importance of mutation rate driving selective sweeps vs. small shifts at many loci of small effect is observed in a model of adaptation in a single population (Höllinger *et al*. 2019, 2023).

Our theory successfully predicts the threshold mutation rate above which mutation swamping occurs. In the parameter range we studied, it was observed at the per-trait mutation rate of 10^*−*3^–10^*−*2^ (Figure 4). Assuming a typical mutation rate of 10^*−*8^ per base, this phenomenon is relevant for traits with a mutational target of 100kb to 1Mb. Although this target size perhaps seems too large for mutation swamping to be associated with many traits, Sella and Barton (2019) argued that the mutational target of many complex traits is around 1 Mb, suggesting that mutation swamping can occur in these traits. Indeed, these complex traits are typically determined by a large number of causal alleles with small effect (Stranger *et al*. 2011; Sella and Barton 2019; Barghi *et al*. 2020), consistent with our prediction.

The threshold mutation rate is lower when the size of the source population is large, the effect size of a new mutation has a large variance, and selection is weak (Figure 4, S6). Although precise estimates of these parameters are still difficult to obtain, and their magnitudes would be expected to vary in more realistic configurations of environment and population structure, we expect that traits would be subject to mutational swamping in species satisfying these conditions. The range of migration rates within which mutation swamping occurs shifts toward lower values as the effect size at the focal locus decreases (Figure 4D). This pattern suggests that mutation swamping will also re-shape the architecture of adaptation through homogenizing small effect alleles while preserving divergence at large effect loci. This effect is analogous to the evolution of a concentrated architecture by migration swamping (Yeaman and Whitlock 2011; Yeaman 2015; Rafajlović *et al*. 2016), although this would occur over lower migration rates in the presence of mutation swamping.

We constructed a theoretical framework based on the diffusion method to consider the joint evolution of a large effect locus and many small effect loci. Although each type of locus has been well analyzed in different frameworks of population genetics and quantitative genetics, their interactions have been rarely investigated (but see Dekens *et al*. 2022). Assuming weak linkage effects, we successfully described their evolutionary interactions. Of note, the mutation rate is an explicit parameter in the present framework, in contrast to the quantitative genetics framework which uses the amount of genetic variance as an indirect measure of mutation rate (Falconer and Mackay 1983). Although the amount of genetic variation does not appear in our theory in the main text, we show that our theoretical framework can give a good prediction for the amount of genetic variation as a function of the mutation rate (see APPENDIX C). In the presence of a diverged locus, migration maintains some genetic variation in population 2 even at a very small mutation rate, which contrasts with the one-population model where the amount of genetic variation vanishes as the mutation rate goes to 0 (Kimura 1965; Turelli 1984; Bürger *et al*. 1989). These findings are therefore consistent with previous simulations (McDonald and Yeaman 2018), which found that genetic variance peaked at intermediate migration rates.

A potential theoretical limitation of our approach is the assumption of linkage equilibrium. Linkage can affect the evolutionary dynamics in the following two ways: reduced effective migration and the Bulmer effect. In the face of gene flow, migrants have an allele set different from residents, and non-small linkage equilibria sustains for several generations even under free recombination. Under divergent selection, this linkage reduces the effective migration rate across the genome. This effect typically arises when fitness of migrants is < *e*^*−*1^ *≈*0.37 (Sachdeva 2022; Zwaenepoel *et al*. 2023), which does not apply in our parameter space. However, if we assume stronger selection, this effect may act and shift higher the range of migration rate within which the transition in architecture occurs. Another potential factor is the Bulmer effect: a negative correlation between the genotypic value at linked loci generated by stabilizing selection on a phenotypic trait (Bulmer 1971). Under this effect, the impact of large effect alleles is essentially reduced because positive effect alleles tend to be linked with negative effect alleles, and vice versa, reducing the stability of the concentrated architecture. This mechanism does not seem to work significantly in the parameter space in this study as our linkage equilibrium approximation explains well the phenotypic variance in simulation (Figure C1). However, if we assume stronger selection or tighter linkage, the Bulmer effect may affect the evolutionary dynamics, deviating it from theoretical prediction.

Our prediction about genetic architecture can be tested by examining the repeatability of the genetic architecture across populations or species. Although it is often impractical to study a time-course of evolution for long enough to establish whether an architecture is stable or transient, we can assume that genetic architecture is stable when the same architecture is observed in comparisons of closely related lineages after a sufficiently long period of independent evolution. Thus, if the same genes repeatedly contribute to the divergence of locally adaptive trait in multiple lineages, we can conclude that the architecture must not be transient, at least at the loci with repeated involvement in adaptation. Based on the behaviour of our model, such traits should have a relatively small mutational target so that the per-trait mutation rate falls below the mutation swamping threshold. By comparing these observations with inferences of the size of mutational target based on GWAS studies (e.g. Simons *et al*. (2022) Figure 3C), we may be able to test the relationship between inferred mutational target size and the kind of architecture that evolves. It should be noted a lack of repeatability does not necessarily mean the action of mutation swamping. In addition to transient architecture, this pattern may also result from diffuse architecture, in which repeatability is low due to the independent evolution under limited migration (see Figure 2A).

Considerable repeatability has indeed been observed in the architecture of local adaptation for many traits (Conte *et al*. 2012; Bomblies and Peichel 2022; Bohutínská and Peichel 2024). The threespine stickleback presents one of the most iconic examples, with repeated adaptation to freshwater environments characterized by suites of genomic islands of differentiation found repeatedly in many populations (Hohenlohe *et al*. 2010; Magalhaes *et al*. 2021; Roberts Kingman *et al*. 2021). While there are few direct estimates of migration rate between marine and freshwater populations, it is commonly assumed to fall within the range of 0.0001 < *m* < 0.1 (Pedersen *et al*. 2017; Galloway *et al*. 2020), which broadly overlaps with the migration rates that would permit mutation swamping. Thus, the observed repeated adaptation implies that the stickleback case falls below the mutation swamping threshold, which suggests a relatively small mutational target size for the freshwater ecotype. However, one might think that it should hardly be the case because the freshwater phenotype involves many characters including feeding, body shape, behaviour, swimming, pigmentation, and body size (Peichel and Marques 2017), which may constitute large mutational target in total. One possibility is that each character evolves in a genetically independent manner and has a small mutational target. Another possible explanation is pleiotropic effect of causal alleles, which is observed in the stickleback case (Archambeault *et al*. 2020) and also in other species (Sella and Barton 2019; Mackay and Anholt 2024). When each causal allele affects multiple traits, the mutational target size involving each trait could be large, but it is also likely to cause deleterious side effects and reduce the effective mutation rate that contributes to the phenotypic variation. Although evaluating this effect is beyond the scope of this study, it would be interesting in future studies to investigate how pleiotropy in multidimensional complex traits affects the genetic architecture underlying local adaptation.

## Data Availability

The authors state that all data necessary for confirming the conclusions presented in the manuscript are represented fully within the manuscript. Codes used for numerical analyses and simulations are available at https://github.com/TSakamoto-evo/polygenic_local_adaptation.

## Acknowledgments

The computing resource was provided by the Digital Research Alliance of Canada and Human Genome Center (the Univ. of Tokyo).

## Appendices Appendix A: The condition for the stable divergence at the focal locus

In this section, we determine the condition that Equation 3 has a stable equilibrium at 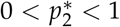. To do so, we first arrange Equation 3 into:

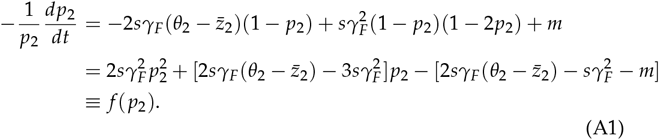

Using *f* (*x*), the condition that 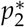 should satisfy is described as:

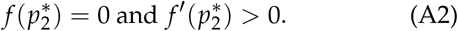

Noting that *f* (1) = *m* > 0, the possible shape of *f* (*x*) is listed in Figure A1. Among these patterns, a stable 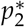 exists in the second and the fifth pattern. Let *x*_axis_ and *D* be the location of the axis of symmetry and the discriminant of *f* (*x*), respectively, where

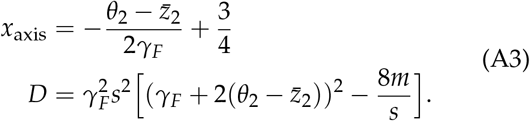

The condition for the second pattern is that *f* (0) *<* 0, which is arranged into:

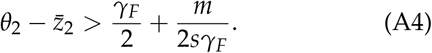

The condition for the fifth pattern is given by:

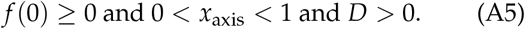

After some calculation, the condition is rearranged into:

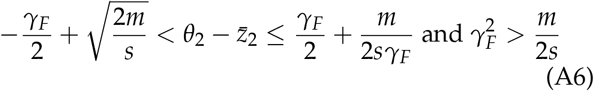

By combining Equations A4 and A6, Equation 4 is derived.

## Appendix B: Derivation of sojourn time based on diffusion method

In this section, we derive the sojourn time density at frequency *x* when evolution starts from a single mutant allele, *τ*_1_(*x, γ*) and 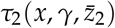.

We first consider the sojourn time in population 1, *τ*_1_(*x, γ*). As we mentioned in the main text, we are specifically interested in the fate of alleles that finally go extinct. Then, we here derive the sojourn time density conditional on the eventual extinction of the mutant allele. Following Ewens (2004), *τ*_1_(*x, γ*) is given by:

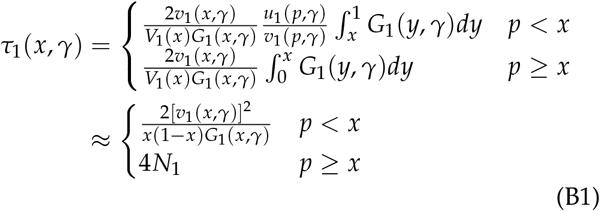

where 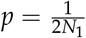 and

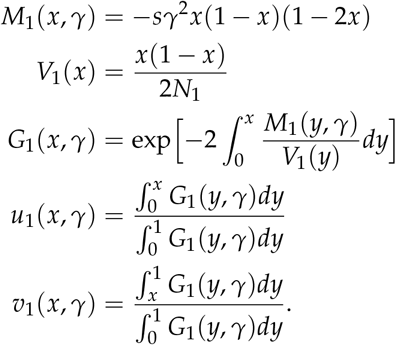

In the approximations, we used that for 0 *< x ≤ p ≪* 1,

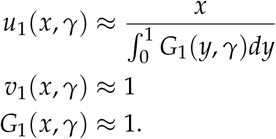

In the main text, *τ*_1_(*x, γ*) is used to calculate 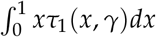 For this purpose, we can approximate

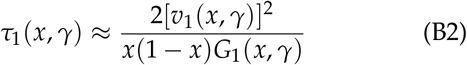

for the entire range of *x* because

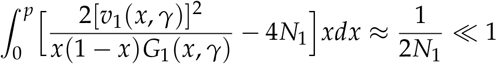

We next consider the sojourn time in population 2. Following Ewens (2004), 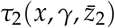 is given by:

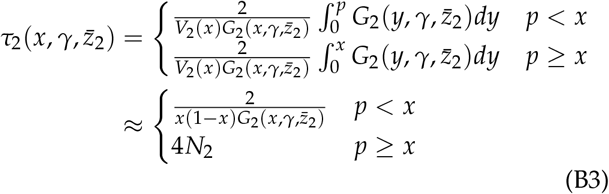

where 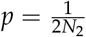 and

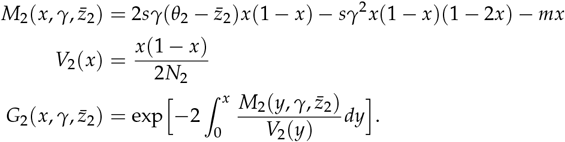

We use 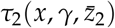 to calculate 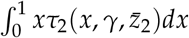 so we can simply assume that

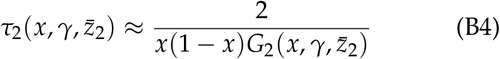

because

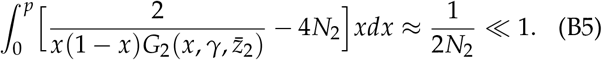

Equations B2 and B4 are used to calculate the numerical results presented in this paper.

**Figure A1.**
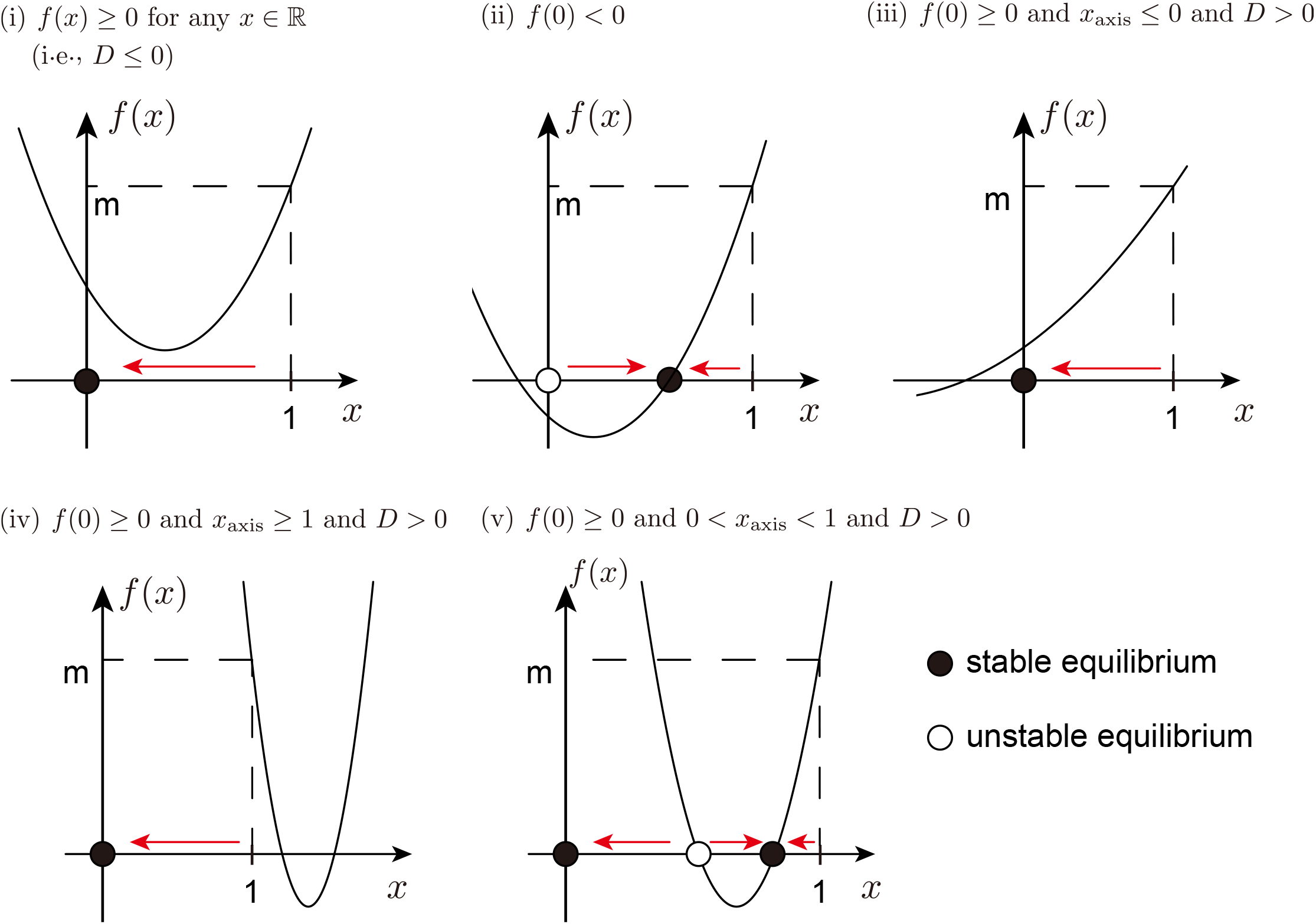
Possible shape of *f* (*x*). *x*_axis_ and *D* is the location of the axis of the symmetry and the discriminant of *f* (*x*), respectively (see Equation A3). Resulting equilibria of *p*_2_ are also plotted on the *x*-axis.

## Appendix C: The amount of genetic variance

The amount of genetic variation (*V*) is a fundamental parameter in a quantitative genetics framework as an indirect measure of mutation rate. Although *V* does not appear in our theory, it would be beneficial to investigate its dynamics for completeness. In this section, we provide an approximation to calculate the amount of genetic variation in population, *V*_*i*_(*µ*), and compare its result with the simulation results.

In the main text, we focused on the contribution of each mutant to the mean genotypic value. Here, we instead focus on the contribution of each mutant to the amount of genetic variation, *V*_*i*_. We first consider the amount of genetic variation in population 1. Since population 1 evolves independently from population 2, it is essentially one population model and previous theories may be applied (Bürger *et al*. 1989). However, for completeness, we present a derivation based on our framework. The total contribution of a mutant allele that arises in population 1 to *V*_1_ can be calculated as:

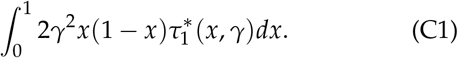

where 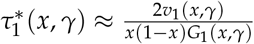 is an expected sojourn time of a new mutation with effect size *γ* at frequency *x*. Note that 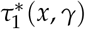 accounts the contribution of a mutation that eventually is fixed and takes a slightly different form from *τ*_1_(*x, γ*), which considers only mutations that are finally extinct. Summing up the contributions from all mutations arising in one generation, *V*_1_(*µ*) is given by:

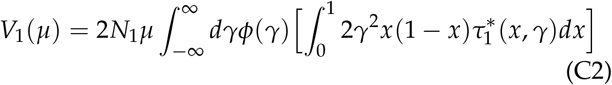

Next, we consider genetic variation in population 2. In the same manner as Equation 6, the total contribution of a mutant allele that arises in population 2 to *V*_2_ can be calculated as:

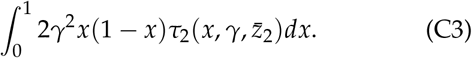

Then, similar to Equation 8, by considering mutations arising in both populations, *V*_2_(*µ*) is given by:

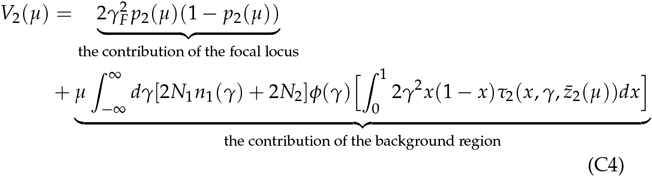

where 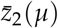 and *p*_2_(*µ*) are the values at equilibrium, which is a function of mutation rate.

In Figure C1, we compare our theory (Equations C2 and C4) with the simulation results. They generally show a good agreement, suggesting the linkage equilibrium approximation is also useful to predict genetic variance. The amount of genetic variation in population 2 shows a complex pattern as a function of mutation rate and migration rate.

**Figure C1.**
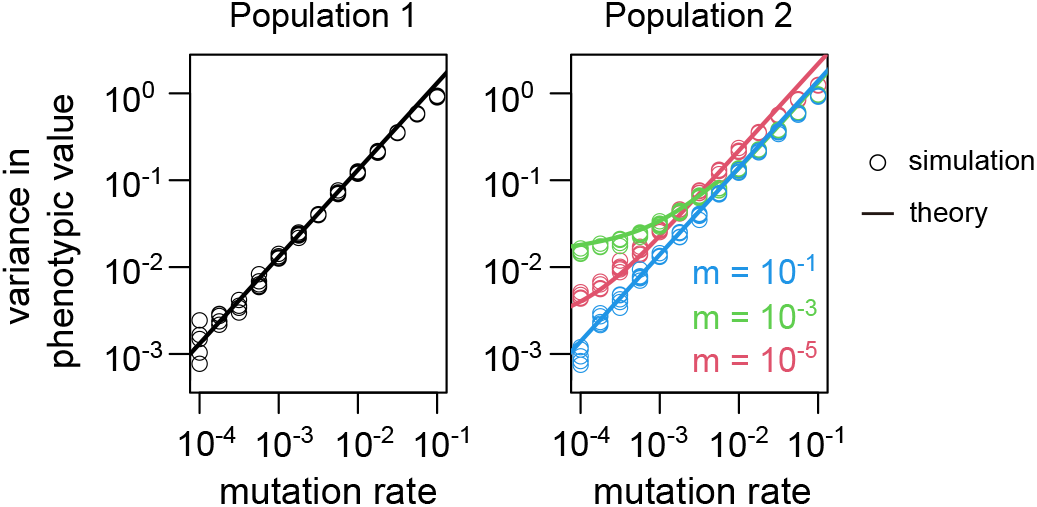
The amount of genetic variation as a function of mutation rate. Parameters *N*_1_ = *N*_2_ = 10, 000, *s* = 0.025, *σ* = 0.02, and *γ*_*F*_ = 0.2 were assumed.

**Table S1.** List of parameters

| parameters | explanation | default | other values investigated |
| --- | --- | --- | --- |
| $m$ | migration rate from population 1 to population 2 | $10^{-5}$–$10^{-1}$ | – |
| $\mu$ | total mutation rate in the background region | $10^{-4}$–$10^{-1}$ | – |
| $\theta_i$ | phenotypic optimum in population $i$ | $\theta_0 = 0, \theta_1 = 1$ | – |
| $N_i$ | size of population $i$ | 10,000 | 50,000 |
| $\sigma$ | standard deviation of effect size at the background region | 0.02 | 0.01, 0.04 |
| $s$ | selection strength | 0.025 | 0.1, 0.25 |
| $\gamma_F$ | effect size at the focal locus | 0.2 | 0.1, 0.4 |

**Figure S1.**
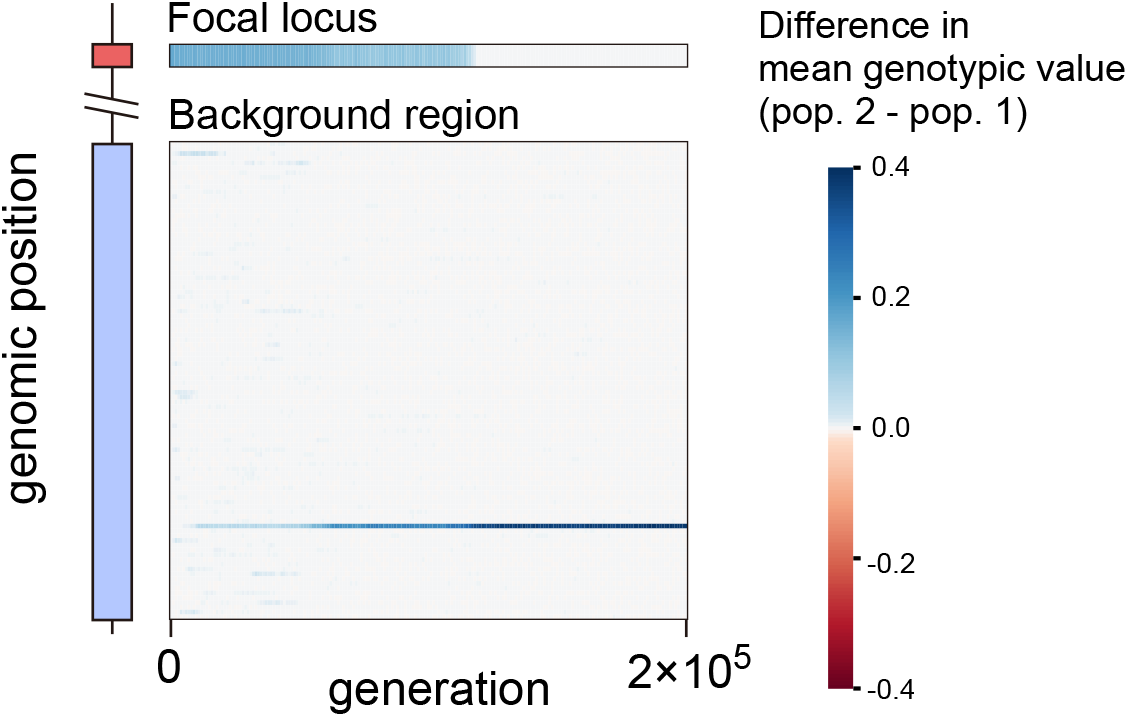
A typical simulation run in which the turnover of the large effect cluster occurs. In this simulation run, *m* = 10^*−*2^, *µ* = 10^*−*2.75^, *N*_1_ = *N*_2_ = 10, 000, *s* = 0.25, *σ* = 0.02, and *γ*_*F*_ = 0.2 were assumed. For visibility, the background region consists of 100 rows, each row representing the sum of the genotypic value difference of 10 adjacent loci.

**Figure S2.**
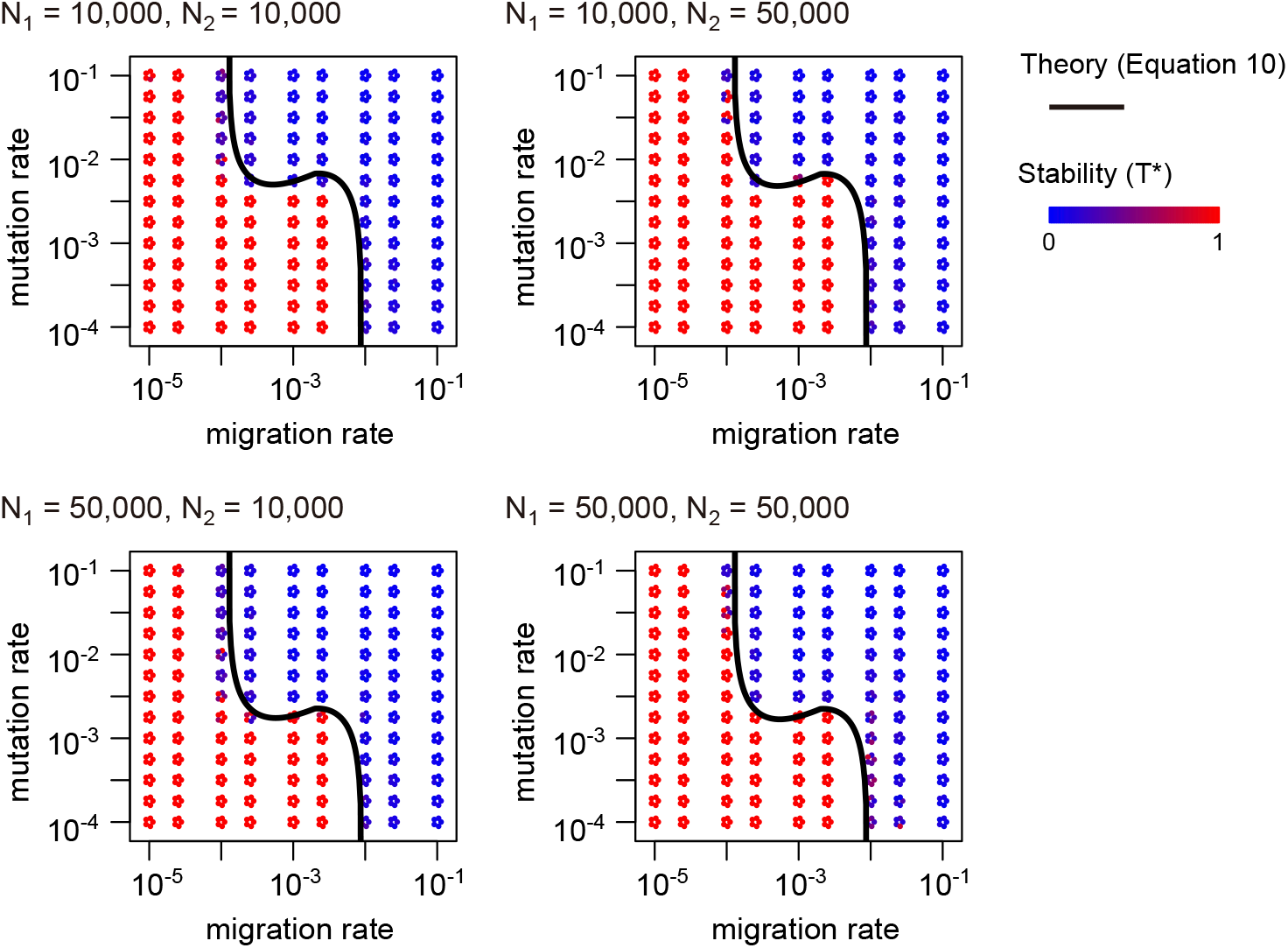
Effect of population sizes on the stability of a genetic architecture in simulation. *s* = 0.025, *σ* = 0.02, and *γ*_*F*_ = 0.2 were assumed.

**Figure S3.**
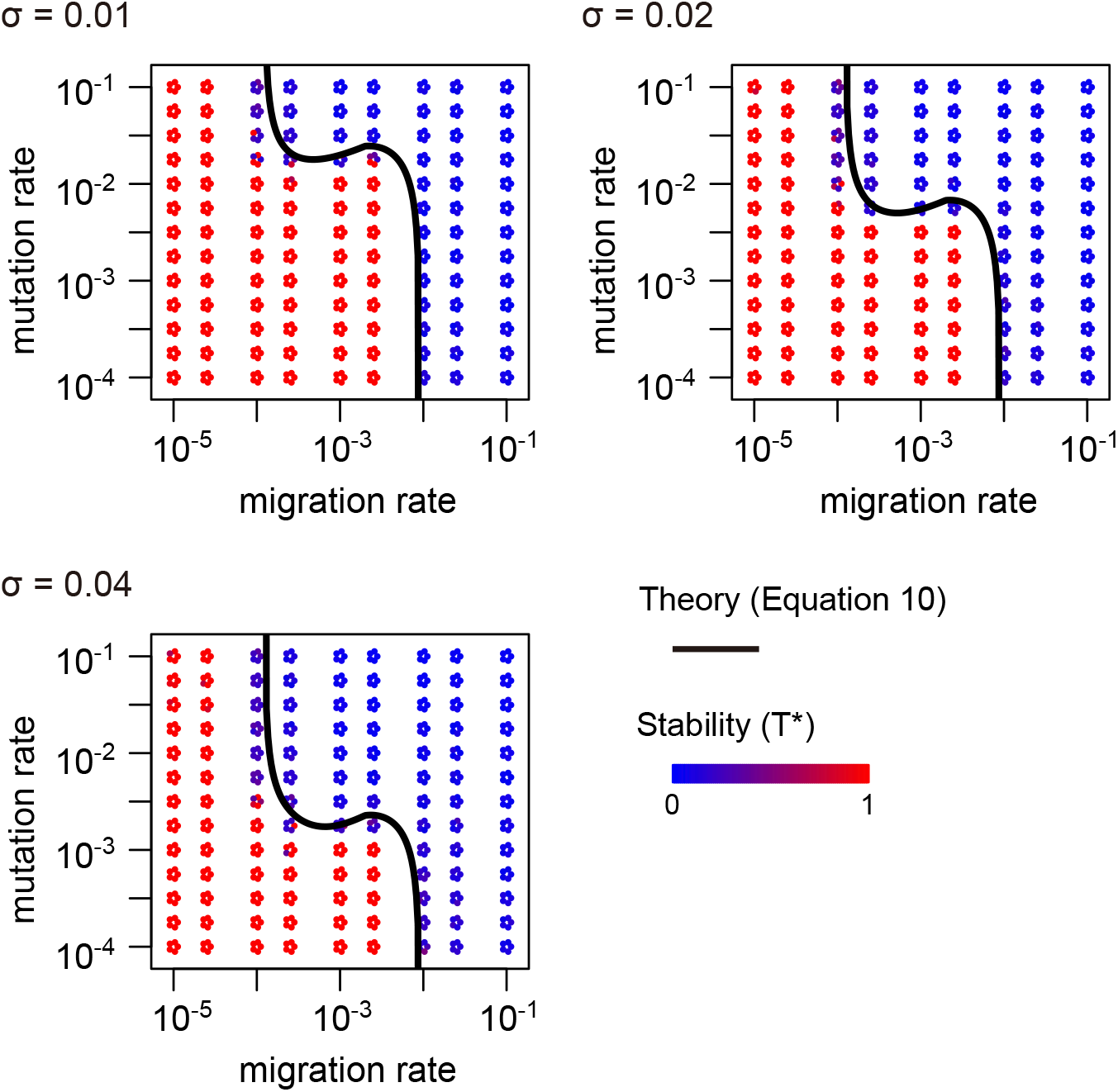
Effect of variation of mutation effect size on the stability of a genetic architecture in simulation. *N*_1_ = *N*_2_ = 10, 000, *s* = 0.025, and *γ*_*F*_ = 0.2 were assumed.

**Figure S4.**
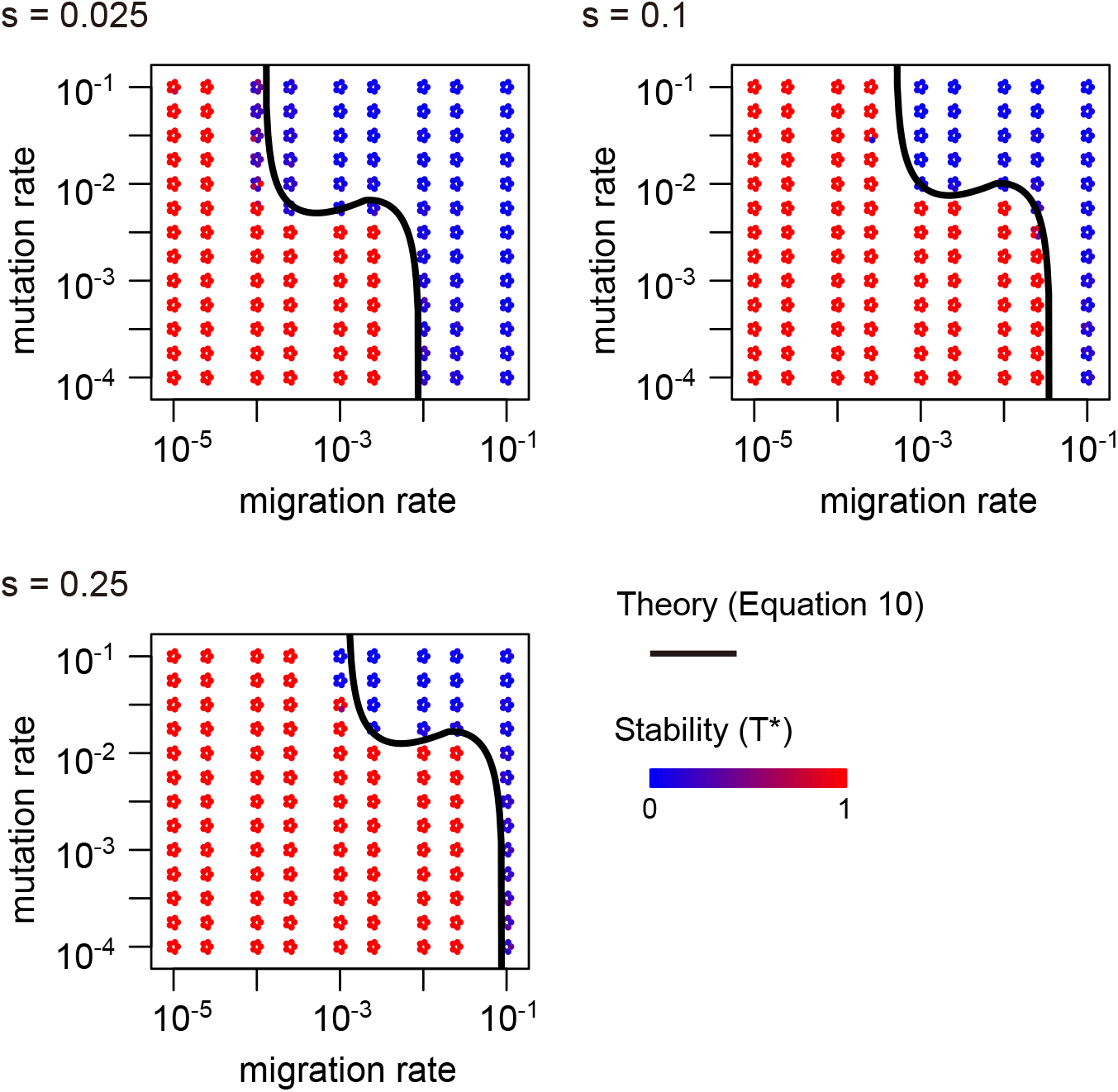
Effect of selection strength on the stability of a genetic architecture in simulation. *N*_1_ = *N*_2_ = 10, 000, *σ* = 0.02, and *γ*_*F*_ = 0.2 were assumed.

**Figure S5.**
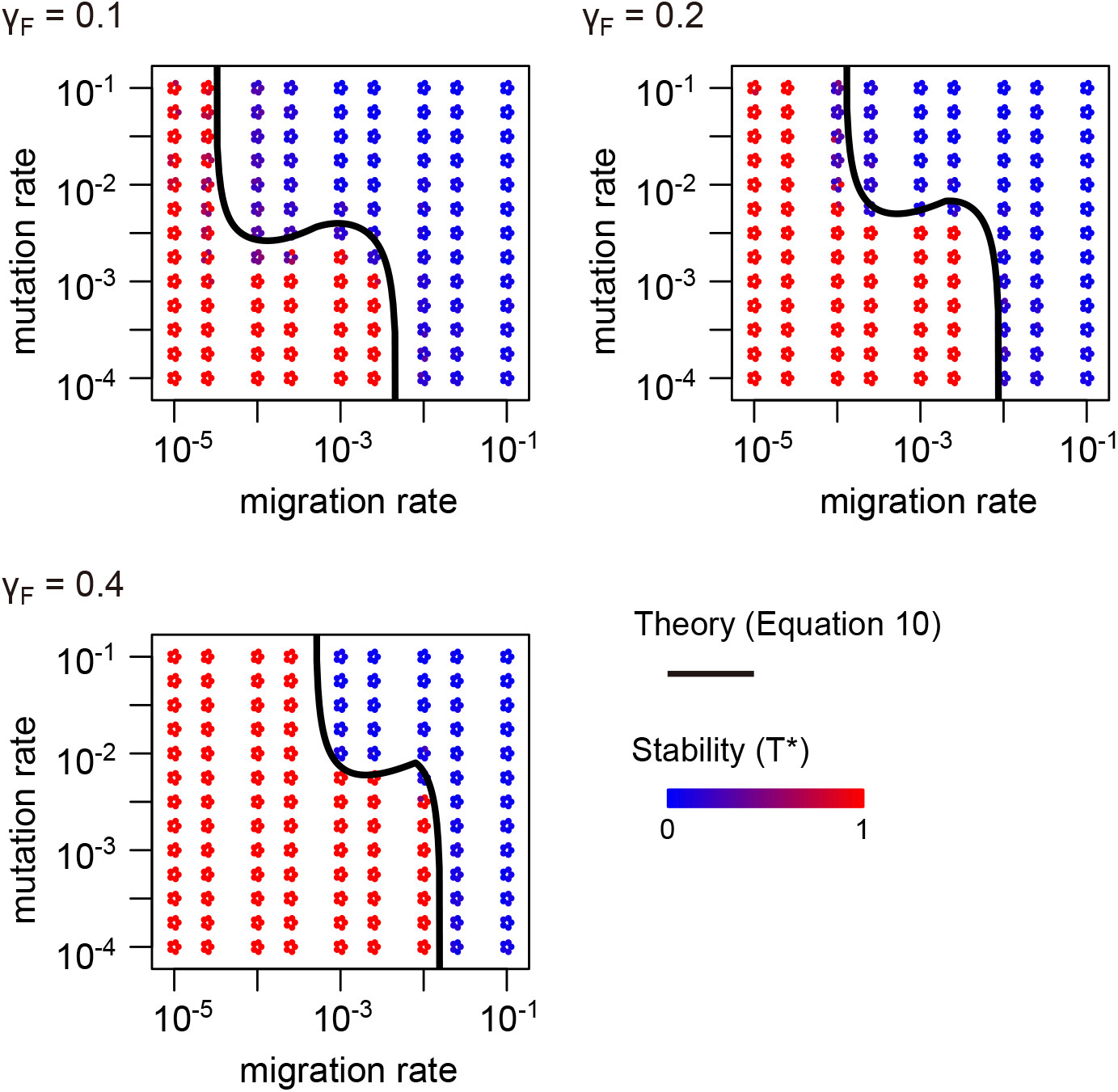
Effect of mutation effect size at the focal locus on the stability of a genetic architecture in simulation. *N*_1_ = *N*_2_ = 10, 000, *s* = 0.025, and *σ* = 0.02 were assumed.

**Figure S6.**
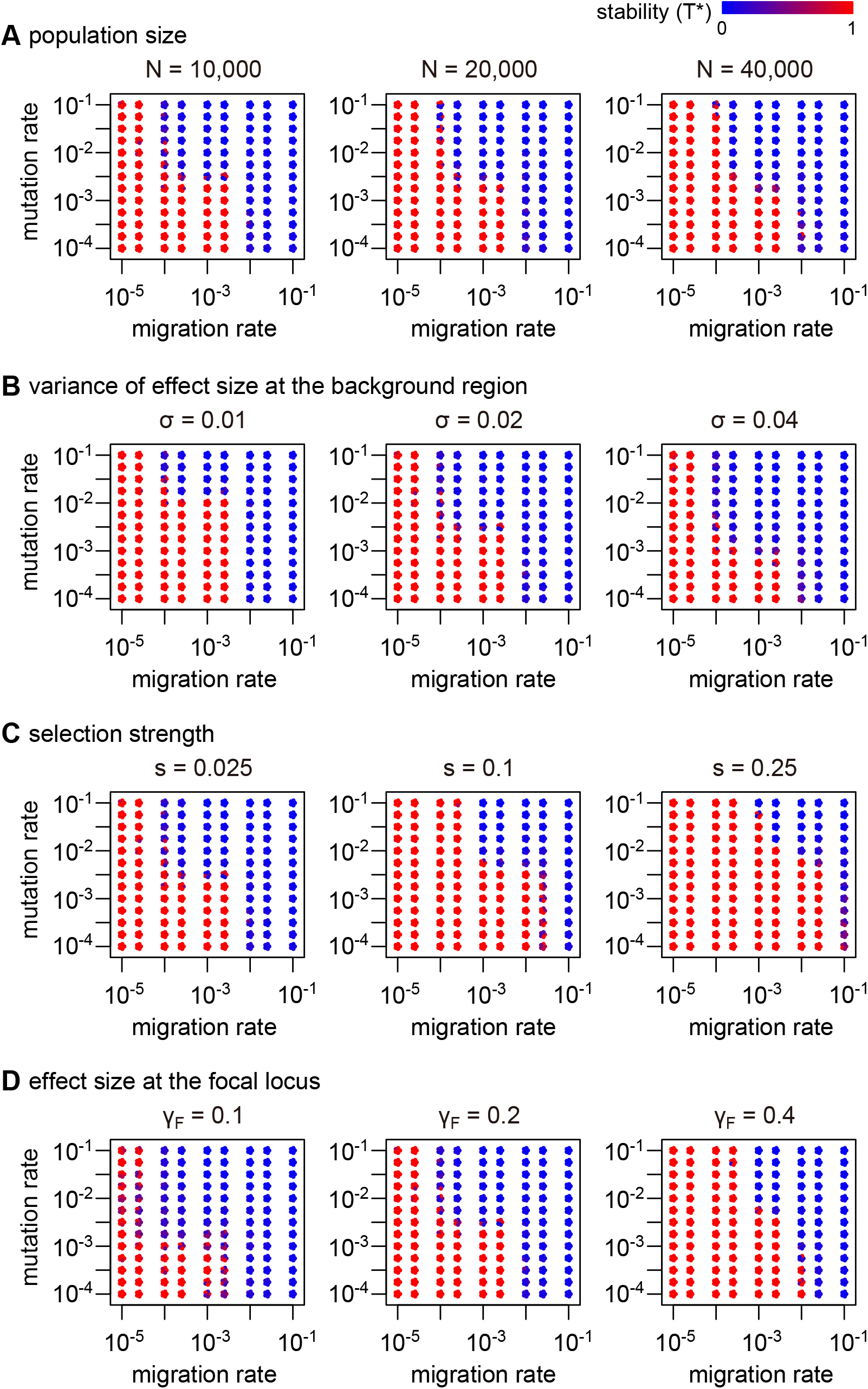
Transition to transience architecture in the symmetric migration model. Unless otherwise specified, *N* = *N*_1_ = *N*_2_ = 10, 000, *s* = 0.025, *σ* = 0.02, and *γ*_*F*_ = 0.2 were assumed.

